# Student-focused development of a next-generation centrifuge force microscope

**DOI:** 10.1101/2020.08.30.274373

**Authors:** K.J. Tompkins, N. Venkatesh, E.T. Berscheid, A.J. Adamek, A.P. Beckman, M.A. Esler, A.C. Evans, B.A. Everett, M. Houtti, H. Koo, L.A. Litzau, A.T. Nelson, T.M. Peterson, T.A. Reid, R.L. Evans, W.R. Gordon

## Abstract

Advanced biological molecule force probing methods such as atomic force microscopy and optical tweezers used to quantify forces at the single-molecule level are expensive and require extensive training and technical knowledge. However, the technologies underlying a centrifuge force microscope (CFM) are relatively straight forward, allowing for construction by labs with relatively low budgets and minimal training. Design ideas from previously constructed CFMs served as a guide in the development of this CFM. There were two primary goals: first, to develop an inexpensive, functional CFM using off-the-shelf and 3D printed parts; and second, to do so in the context of providing an educational experience for a broad range of students. The team included high school students and undergraduates from local high schools, the University of Minnesota, and other local higher education institutions. This project created an environment for student-focused development of the CFM that fostered active learning, individual ownership, as well as excellence in research. The instrument discussed herein represents a fully functional CFM designed and built by a postdoctoral researcher and a graduate student who together mentored several high school and undergraduate students.

**STATEMENT OF SIGNIFICANCE:** The presented centrifuge force microscope (CFM) builds on features of existing designs specifically engineered for probing macromolecular force interactions at the single-molecule level. In the coming years, more versatile and modular CFM designs will be utilized in the force spectroscopy field, and the presented design is a step in that direction. In addition to constructing a functional instrument, true student ownership of the project design was equally an end goal. Students from high school through graduate school were included, and the project was structured so that everyone was seen as peers. This active learning project allowed students to acquire scientific concepts and techniques and apply them to real-life situations.

## INTRODUCTION

### History of Centrifuge Force Microscopy

Biomechanical forces manipulating interactions between, or within, macromolecules are of great interest to the scientific community, from understanding cancer metastasis progression (Chaudhuri et al., 2014; Ma et al., 2018; Shieh, 2011; Sottnik et al., 2015) to quantifying mechanical stability of individual proteins (Borgia et al., 2008; Le et al., 2018; Lv et al., 2014; Rief et al., 1999, 2000; Schlierf et al., 2004; Sharma et al., 2007). Single-molecule force measurements can provide information on the strength of interactions between individual molecules, and these experiments primarily use force probing instruments such as atomic force microscopy (AFM), optical tweezers (OT), and magnetic tweezers (MT) (Neuman & Nagy, 2008). However, AFM and OT can be especially expensive and require a high level of user expertise. Particularly at smaller institutions, the cost of the instrumentation and supporting infrastructure, in addition to the required expertise, impedes the implementation of these technologies.

CFMs were introduced as early as 1930 when E. Newton Harvey and Alfred L. Loomis published their design of a hand-cranked CFM for the study of living cells (Harvey & Loomis, 1930). Since then, these instruments have been adapted for the study of many aspects of biology, from cytoplasmic streaming and cell motility (Fukui et al., 2000; Hiramoto & Kamitsubo, 1995; Kamitsubo et al., 1989) to more recent designs capable of quantifying single-molecule biomolecule rupture forces (Halvorsen & Wong, 2010; T. Hoang et al., 2016; T. P. Hoang et al., 2015).

These later designs apply forces on the protein(s) of interest, which are tethered to glass on one end and a bead on the other, by spinning a microscope focused on the experimental chamber. (see Figure 1). The applied force is equivalent to the centripetal force needed to pull the tethered bead in a circular path. The presented CFM can tether hundreds or even thousands of molecules/molecular complexes of interest to the experimental chamber’s glass surface simultaneously, allowing for experimental multiplicity. Rupture events are recorded as single beads leave the focal plane at distinct timepoints under an applied force. In the described validation experiments, the rupture point occurs at the covalent DIG:α-DIG interaction because it is weaker than the streptavidin:biotin interaction (see Figure 5). The DIG:α-DIG is utilized because it is a popular site-specific anchor point for protein-DNA conjugation in force spectroscopy due to ease of assembly and specificity. The value of the force (*F*) is calculated as shown in equation 1, where the force is the product of the buoyancy adjusted bead mass (*m*), the square of the angular velocity (ω), and the radius of the circular path (*r*).

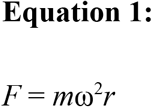

**Figure 1:**
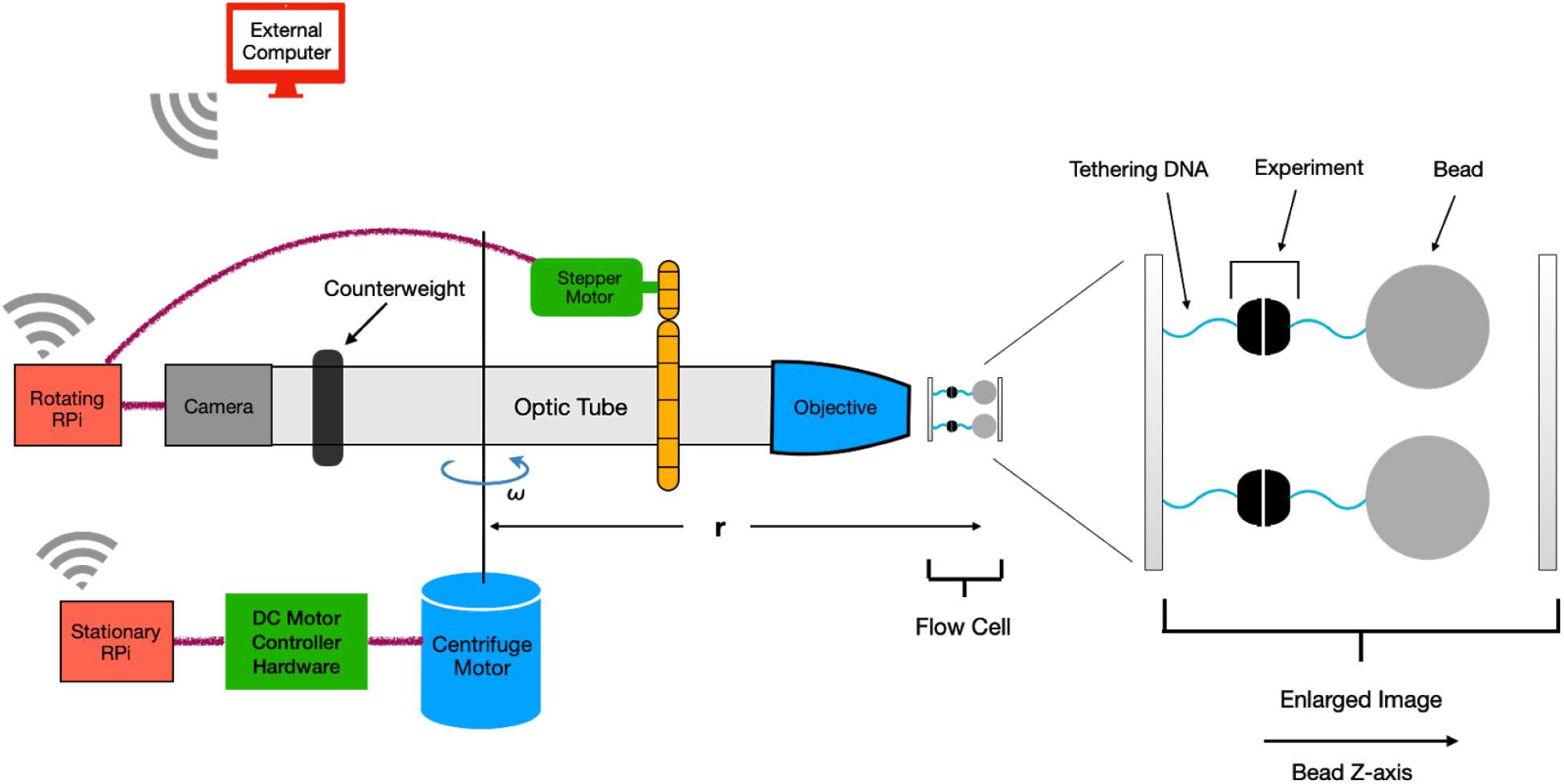
Schematic of CFM design with experimental sample. The figure depicts the overall design and layout of the CFM including an example of a biological sample in the experimental chamber. Further detail regarding the design is provided in the Materials and Methods section.

### Educational Emphasis

This student-centered project brought together a team of high school, undergraduate, and graduate students, who were managed by a graduate student and a postdoctoral researcher who sought to fulfill teaching and research purposes through the development of a working CFM. This project incorporated biomechanical engineering, biophysics, and biochemistry, providing student researchers with multiple learning opportunities. For example, students learned and utilized Python programming skills, Linux OS interface skills, design engineering (electrical and mechanical), and biochemistry techniques, while advanced researchers gained multi-disciplinary management experience. As a team, we successfully designed and constructed a working CFM, based on previously published designs, and briefly validated force-dependent rupture events of the digoxigenin:anti-digoxigenin antibody (DIG:α-DIG) interaction using the instrument.

From start to finish, the educational practice known as ‘active learning’ was employed (cite). The collaborative laboratory/design project was an ideal environment for satisfying all three learning domains considered to be essential to deep, applied understanding (Bransford et al. (1999)). For the ‘cognitive domain,’ the project provided new information, such as an understanding of both biological and engineering systems and methods with their application to concrete goals (Bransford). A wide breadth of skills, from machining and 3D design to pipetting and lab safety, directly addressed the ‘psychomotor domain’ (Bransford). The third learning domain, the ‘affective domain’ (Bransford), was addressed by student integration into leadership roles, which reinforced their internal motivation and sense of value to the team. These concepts were promoted through the belief that student participation is essential and appreciated, even more than the biology and construction of the instrument. In other words, student involvement in research was considered more important than the research itself.

‘Knowledge-centered’ learning environments place the student first, with the primary goal of helping students become knowledgeable by learning in ways that lead to deep understanding, including subsequent transfer of knowledge (Bruner, 1981). So, in addition to establishing an active learning environment where students acquired knowledge and experience, the students were also positioned to be teachers and leaders. To create learning opportunities for high school and undergraduate students, the various subtasks for the CFM’s design and assembly were delegated to smaller teams. These subtasks consisted of computer software development, centrifuge design, microscope design, electronics, and biological testing (see table 1).

**Table 1:** Delineation of the subtask and teams along with the contributors’ educational levels. Abbreviations: PD = postdoc, GSL = graduate student(s) from this lab (Gordon Lab), GSO = collaborating graduate student(s) from other lab(s)/department(s), UGU = undergraduate student(s) from the University of Minnesota, UGO = undergraduate student(s) from other institution(s), HS = high school student(s).

| Subtasks | Team Responsibilities | Educational Levels Represented in the Team |
| --- | --- | --- |
| Project Management | • Overall Leadership<br>• Teams Leadership | → PD<br>→ See each team |
| Computer | • RPi-based design:<br>Leadership<br>Membership<br>• RPi Linux installation and security:<br>Leadership<br>Membership<br>• RPi GIT Hub software version maintenance:<br>Leadership<br>Membership<br>• RPi image capture software:<br>Leadership<br>Membership<br>• Centrifuge control software:<br>Leadership<br>Membership<br>• User interface software:<br>Leadership<br>Membership | → PD and GSL<br>→ PD and GSL<br>→ UGU<br>→ UGU<br>→ UGU<br>→ UGU<br>→ PD and UGU<br>→ HS and UGU<br>→ HS/UGU*<br>→ HS/UGU* and HS<br>→ PD and HS/UGU*<br>→ PD and HS/UGU* |
| Electronics | • All (dual computer control, power supply, centrifuge motor control, stepper motor control, accelerometer/vibration, braking):<br>Leadership<br>Membership | → HS/UGU*<br>→ HS/UGU* and HS |
| Centrifuge | • Chassis:<br>Leadership<br>Membership<br>• Motor, mount: | → PD and GSL<br>→ PD, GSL, and UGO |
|  | Leadership<br>Membership<br>• Accelerometers (vibration Detection, braking):<br>Leadership<br>Membership | → PD and GSL<br>→ PD and GSL<br>→ HS/UGU*<br>→ HS/UGU* |
| Rotor | • Design and build<br>Leadership<br>Membership | → GSL<br>→ GSL |
| Microscope | • Optics<br>Leadership<br>Membership<br>• Focusing mechanics<br>Leadership<br>Membership | → GSL<br>→ GSL<br>→ PD and GSL<br>→ PD and GSL |
| Biology | • Validation testing<br>Leadership<br>Membership | → GSL<br>→ GSL, UGU, and UGO |
| Documentation and Writing | Leadership<br>Membership | → PD and UGU<br>→ everyone |

\* This member started as a high school student (first summer), but thereafter, continued as an undergraduate student from another institution.

Students were also initially encouraged to explore unfamiliar areas of the project. For example, students with no prior experience in programming were encouraged to sign up for Python programming language course(s) from online sources (Codecademy and SoloLearn). Students who had never picked up a hammer were encouraged to take a machine shop course offered by the university’s Physics Department. Others were encouraged to watch videos or seek out training from peers to become proficient in using selected 3D, computer-aided design software (SketchUp Make) and printer control software (Repeteir-Host and Slic3r). Students began working on smaller projects in different fields/subdisciplines before choosing the subprojects that best aligned with their strengths and interests. Students demonstrating proficiency in leadership, mentoring, and/or teaching skills were often made lead over a subproject. These student leaders then led and trained their team of high school and undergraduate peers.

From a management perspective, one of the most challenging aspects of this education philosophy was accepting student contributions that would work but for which the management had a ‘better idea’. It was more important to integrate a student’s ideas than to optimize or replace it with the manager’s ideas. However, advice and support was still provided in situations where it was needed.

### Centrifuge Force Microscope Design

Recent CFMs designed for single-molecule force spectroscopy use sample chambers and optical components integrated within a swinging-bucket of a commercial centrifuge (T. Hoang et al., 2016; T. P. Hoang et al., 2015; Kirkness & Forde, 2018; Yang et al., 2016). These versions are more easily constructed due to their use of commercially available centrifuges and the availability of shared, 3D printable part files. However, these compact, commercial centrifuge designs have greater volumetric constraints due to the limited size and shape of the swinging buckets. Therefore, the incorporation of components that provide additional capabilities, such as on-board focusing, has not been achieved.

Unlike CFMs that utilize commercial centrifuges, the presented CFM uses a custom-made rotor that spins on a custom-built centrifuge, similar to a CFM design published in 2010 (Halvorsen & Wong, 2010). The goal of this active learning (Michael, J., Where’s the evidence that active learning works? Adv Physiol Educ 30: 159–167, 2006; doi:10.1152/advan.00053 and Lopatto, D., Undergraduate Research Experiences Support Science Career Decisions and Active Learning, CBE—Life Sciences Education Vol. 6, 297–306, 2007) project was to design and construct a working, WiFi-enabled, CFM while placing high emphasis on student engagement. The CFM would be of our own design and would explore new features, such as real-time focusing, real-time image acquisition, and data analysis. The electronics control system was built on the educationally-popular Raspberry Pi (RPi) platform (Raspberry Pi Foundation, Raspberry Pi (Trading) Limited, Cambridge, UK) because many RPi-specific, student-oriented training modules are available for learning Linux, basic electronics, and Python programming.

While static balance could easily be achieved by mounting the rotor/microscope assembly on a horizontal shaft, “dynamic unbalance”—balance problems that only occur while spinning—is difficult to overcome without sophisticated balancing mechanisms (El-Saeidy & Sticher, 2010).

The issue of dynamic unbalance, vibration, and safety were always top priority considerations. To mitigate some of the vibration and safety issues, the presented CFM validation experiments used larger streptavidin-coated beads (4.5 μm diameter), which required lower RPMs to achieve sufficient force ((Table XX of forces and rpms??)). The advantages and disadvantages of using larger versus smaller beads is addressed in the Discussion. Additional ideas for mitigating dynamic spin unbalance in a non-swinging bucket design are described in further detail in Future Directions.

## MATERIALS AND METHODS

This section provides a very brief description of the CFM construction and the experimental setup used for instrument validation. More details, including drawings and descriptions of machined and 3D-printed parts, wiring schematics, custom Python software, and lists of purchased materials, can be found in the supplemental materials.

### Mechanical, Optical Parts, and Electronics

All rotating components were mounted to a 12” x 12” x ¼” (thick), precision-machined, aluminum plate. The plate was bolted to a Swinging-4-bucket rotor (from a Beckman Coulter, Allegra 6KR centrifuge). This plate+swinging bucket rotor assembly is referred to as the ‘CFM rotor’ or ‘rotor.’ Onboard components were attached to the CFM rotor using nuts and bolts of appropriate dimensions. To accommodate this particular Beckman Coulter rotor, which was designed to fit onto a 15 mm shaft, the custom centrifuge utilized a shaft of that size. The shaft was mounted to a DC motor by way of a flexible coupling that prevents vibrational damage to the motor during spinning. The motor and 15 mm shaft assembly were mounted in a custom, wooden frame using heavy-duty radial and thrust bearings. The wooden frame was constructed with plywood and two-by-fours purchased at a local lumberyard. To further dampen vibration in the wooden frame construction, the entire centrifuge was reinforced with heavy-duty steel, C-channel framing. Then the entire CFM was attached on top of a 36” x 36” x 2” (thick) steel plate, the plate having been salvaged from a discarded optical table (see Figure 2). The centrifuge DC motor was controlled by a Raspberry Pi 3 (“stationary RPi”) through a DC motor controller. Detailed wiring diagrams and a parts list are available in supplemental materials.

**Figure 2:**
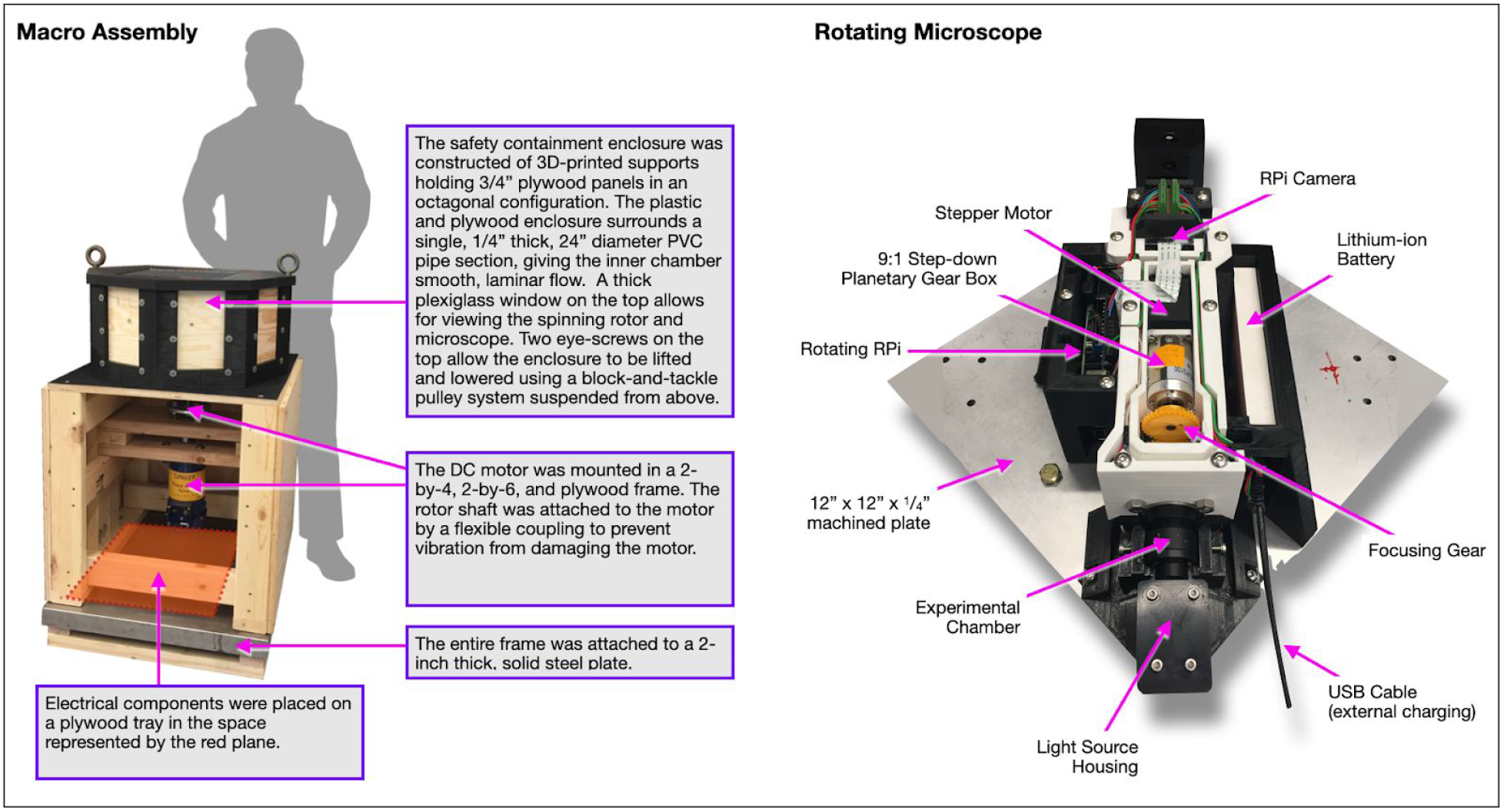
Showing the macro assembly and the rotation microscope on a 12” by 12” plate. The use of a larger, custom rotor design permits the exploration and potential addition of new features to the CFM, a few of which are addressed in Future Directions. However, every design includes compromises, and using a custom-built centrifuge design means that the presented CFM is more complex to build, which reduces reproducibility. To improve reproducibility, the use of proprietary components was minimized, and off-the-shelf parts were preferred. For non-commercially available components, 3D printed parts were first modeled using SketchUp (Trimble Inc., Sunnyvale, CA). Build descriptions for the mechanics, optics, electronics, and controlling computer software are provided as supplemental materials. See supplemental_master_document.pdf, which serves as a map to all of these resources.

The CFM rotor held the following microscope components: an experimental chamber and light source; focusing hardware comprised of a stepper motor, custom gears, and a bearing-mounted, optical shaft that could focus the objective (Olympus xxxxx); various mirrors; a Raspberry Pi 3 (“rotating RPi”) equipped with a stepper hat (Adafruit Industries LLC); and a lithium-ion battery. Figure 1 provides a diagrammatic view of the CFM microscope, and Figure 2 provides a top-down photograph of the CFM microscope.

In order to focus the CFM microscope while spinning, the objective was mounted in a custom-machined optical tube “caged-in” by 8 radial bearings, mounted on bearing cage rods and screwed into the experimental chamber (see Figure 3). These 8 bearings allow the optical tube to turn freely and slide. Guided by live-streamed images captured and transmitted by the rotating RPi, the user focuses by way of a stepper motor, engaging a focus gear that can screw the optical tube (and contained objective) into and out-of the experimental chamber. The focus gear design also allows for a sliding action with the stepper motor gear. This configuration allows the objective to focus while spinning. The optical tube was balanced over the axis of rotation with a counter-weight at one end, providing mass balance for the objective at the other end (see Figures 1 & 3). The distance from the axis of rotation to the experimental chamber was 15 cm. While the movement of the focusing tube/objective did change the balance slightly, increases in vibration were not detectable by an iPhone (Apple, Inc., Cupertino, CA) mounted to the centrifuge to measure vibrational changes using the phone’s accelerometer coupled with force measurement software (Sensor Kinetics Pro, INNOVENTIONS, Inc, Houston TX).

**Figure 3:**
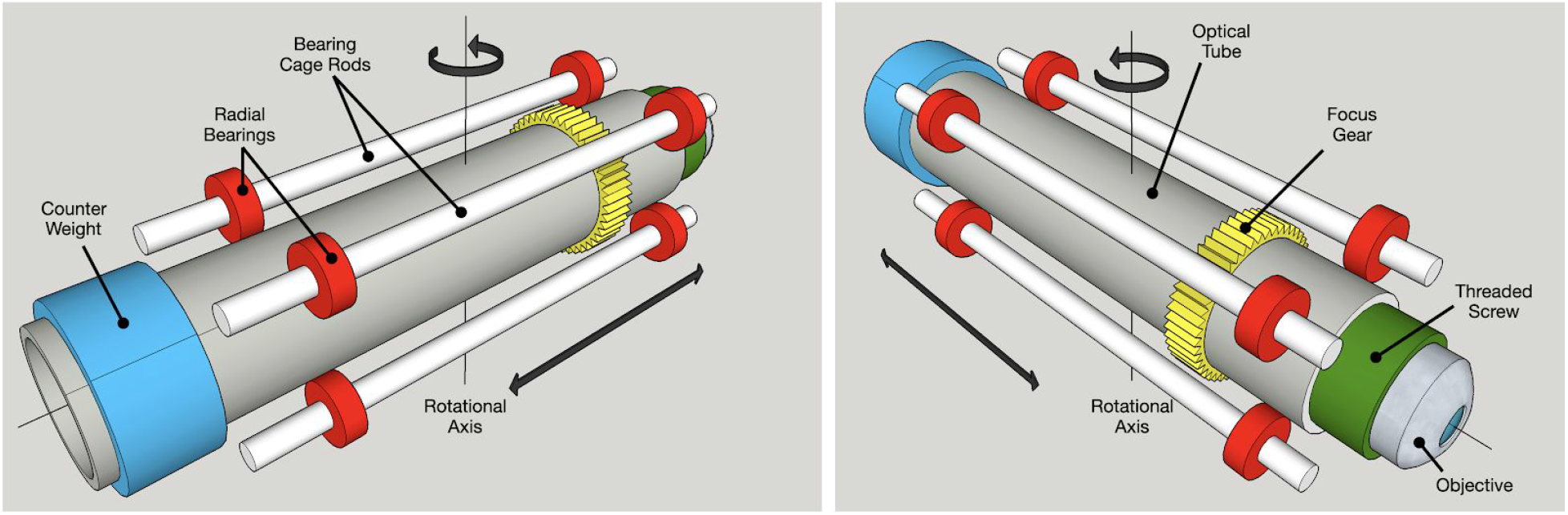
Showing the basic components, from two different angles, supporting real time focusing. The software used to produce this figure (SketchUp Make, Trimble, Inc.) was the same software that, along with Trimble-supplied STL-converter/export extension, was used by students to design components for 3D printing.

### Electronics (Schematics and Function)

The electronics can be divided into two categories, the stationary electronics and the rotating electronics. The stationary electronics in the base of the centrifuge are responsible for controlling the DC motor and includes all of the stationary electrical components. The electronics mounted on the rotor-microscope assembly are referred to as the rotating electronics. Both the stationary and rotating electronics include Raspberry Pi computers (stationary RPi and rotating RPi) that communicate with one another over Wi-Fi (See Figure *4*). The user controls the CFM from an external, Internet-connected computer using a top-level user interface that ‘talks’ only to the rotating RPi. The rotating RPi then sends the desired RPM-time profile to the stationary RPi controlling the DC motor. Once the assembly begins spinning, the rotating RPi captures images and sends them to the external, controlling computer for focusing, bead counting, curve-fit analysis, and data storage. Communication between all three computers is handled via SAMBA (for file transfer) and SSH (for user interface/control). During this whole process, the stationary (motor-controlling) RPi also monitors vibrations in all 3 dimensions by way of an x, y, z-accelerometer component. In the case of abnormally high vibrations, the rotating Pi can cut power to the DC motor and perform an emergency halt via magnetic braking.

**Figure 4:**
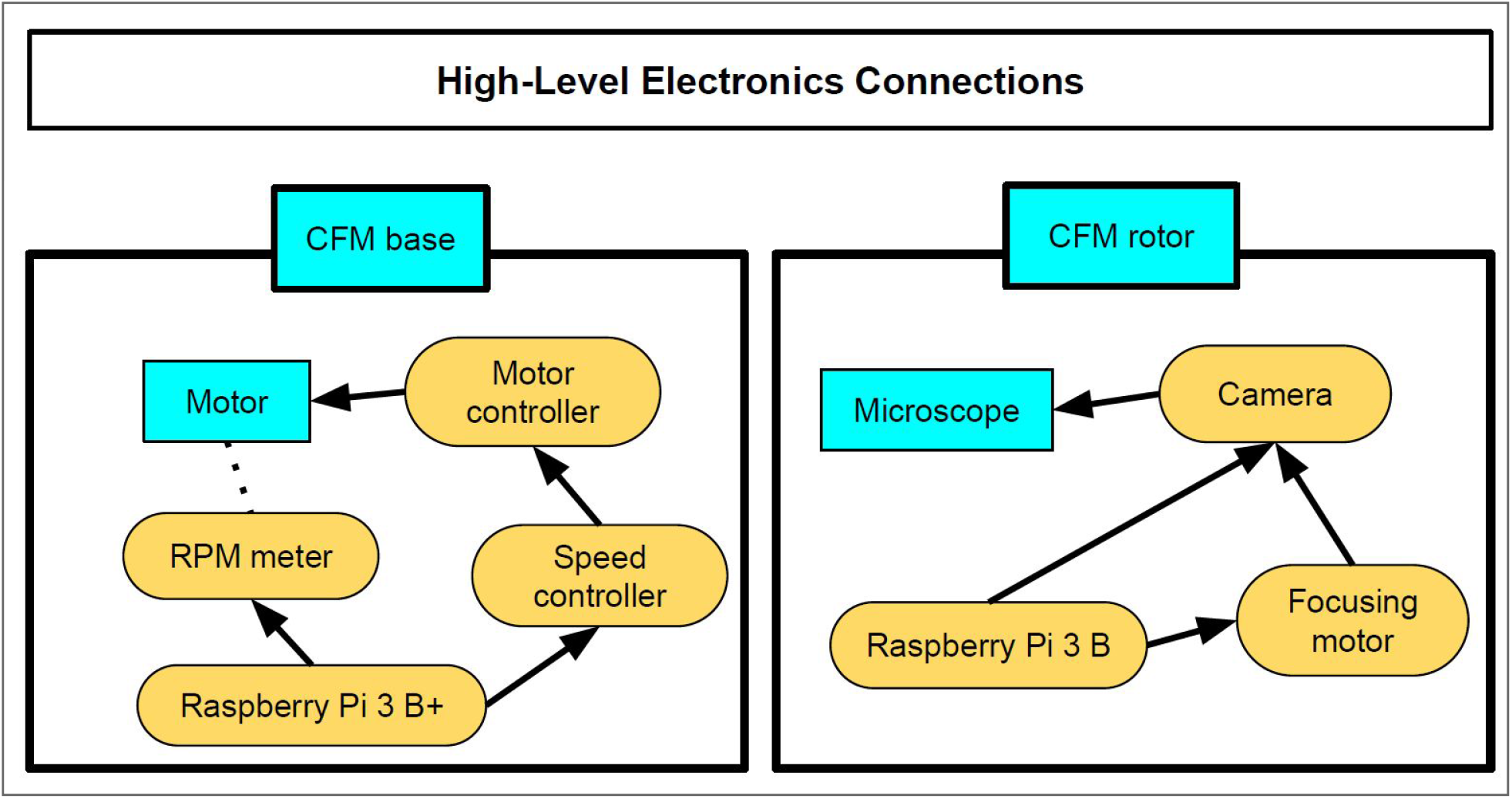
Information and Control Flow. Arrows indicate master/slave relationship; the dotted line represents sensor data; yellow ellipses are electronics modules; blue boxes designate mechanical assemblies.

### Image Capture and Processing

The CFM microscope uses a second-generation, 8 megapixel Raspberry Pi camera module (V2) with the included, fixed focus lens removed so that the Sony Exmor IMX219 sensor of the camera can be connected to the purpose-built microscope. A stationary Raspberry Pi computer controls the centrifuge spin and the camera’s images are captured by the rotating Raspberry Pi. Raspistill software (Raspberry Pi Foundation) is used to acquire time-lapse images at a rate of approximately one image per 5 s. The accuracy of the bead counting software (see XXXXXX.XXX in supplemental materials), written in MatLab, was validated by comparing the calculated counts to manual counts.

### 5’ DIG 3’ Biotin DNA Constructs

To validate the CFM’s function, a flow cell containing tethered 4.5 μm beads (mass: xxx μg, adjusted/in-solution mass: xxx μg) was constructed. The rationale behind the validation testing was not to revalidate previous biological construct(s) used for tethering but rather to validate the presented instrument’s ability to reliably apply precision forces resulting in bead rupture events. Beads were tethered to glass in a flow cell as described by Yang et al (Yang et al., 2016). The tethering DNA was comprised of 132, 60 base pair oligo sequences complexed to a single-stranded M13mp18 DNA handle. This construct is also described in Yang et al. (Yang et al., 2016) and illustrated in Figure 5. Either biotin or digoxigenin (DIG) covalently linked DNA oligos are complemented to either the 3’ or 5’ end of the DNA handle, respectively. A full list of the 60 base pair oligo sequences can be found in supplemental file, S__.

**Figure 5:**
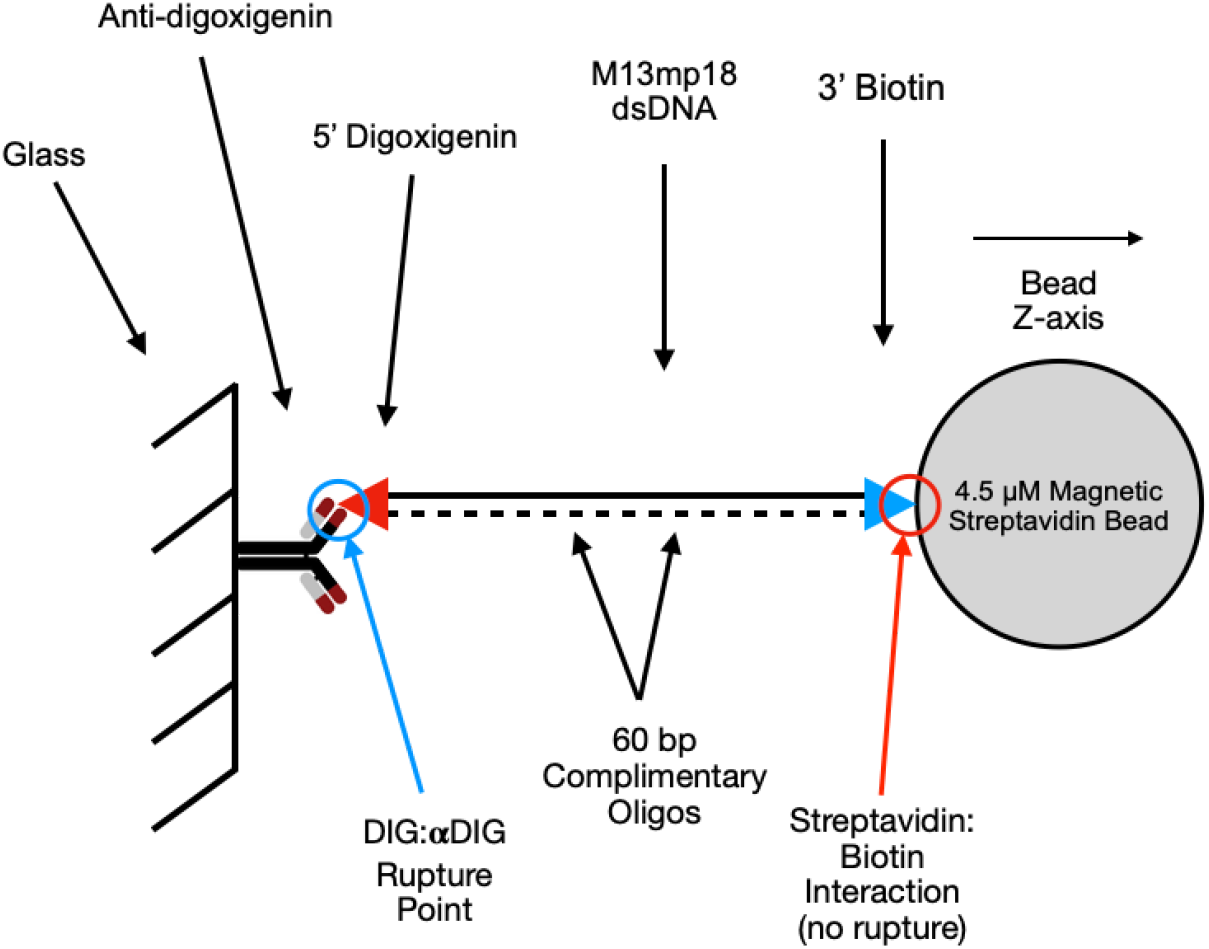
Sample Construct. Depicting the sample construct used to tether streptavidin beads to the glass surface of the flow cell. The covalent interaction of the DIG:αDIG is weaker (XX pN) than the streptavidin:biotin interaction (YY pN). Therefore, when <YY pN of force is applied to the sample construct, the DIG:αDIG interaction ruptures causing the bead to exit the focal plane.

**Figure 6:**
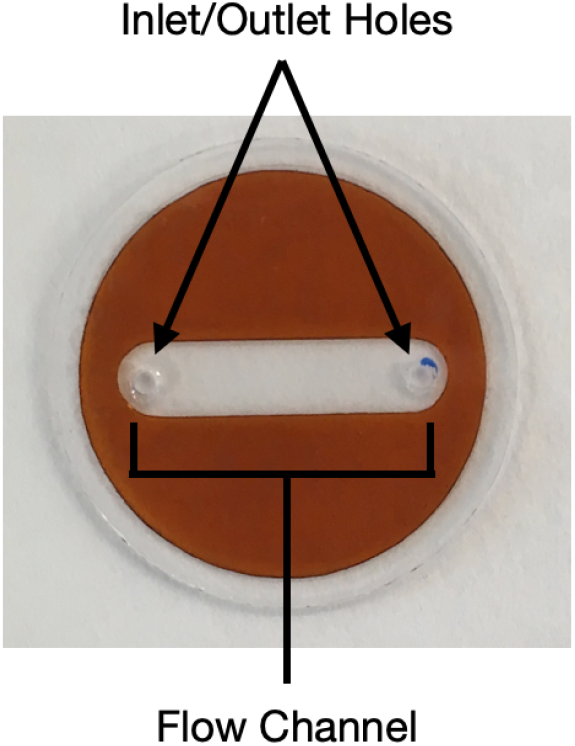
A view of the assembled flow cell showing the bored, inlet and outlet holes and cut flow channel.

### Flow Cell

A flow cell was prepared by sandwiching double-sided polyimide tape (Kapton tape) between a 25 x 25 mm circular support glass (custom cut, Howard Glass, Co.) and a 22 x 22 mm circular cover glass (Hampton). Two, 1 mm holes were drilled into the 0.7-mm-thick support glass as inlet and outlet holes. Prior to assembly, both the support and cover glass were washed by immersion in a near-boiling 2% (v/v) Hellmanex III solution (Sigma-Aldrich) followed by sonication for 30 minutes. The glass pieces were then rinsed with heated Milli-Q water before sonication for another 30 minutes in Milli-Q water. A 20 x 4 mm flow cell was cut from a 22 mm diameter piece of Kapton tape using a Silhouette Cameo® (silhouetteamerica.com). Both pieces of glass were then dried under air flow. A thin layer of 0.2% nitrocellulose in amyl acetate was applied to each slide and baked for 10 min at 80°C before sandwiching the cut, Kapton tape between the nitrocellulose coated, glass slide and the support glass, creating the flow cell’s flow channel. The flow cell volume capacity is ~15 μL.

### Bead-to-glass tethering construct

For flow cell preparation, 20 μL of 5 μg/mL anti-digoxigenin (α-DIG) (Sigma; in PBS pH 7.4) was injected into the flow cell and twice allowed to incubate at room temperature for 15 minutes. The chamber was washed four times with 20 μL SuperBlock (ThermoFisher; in PBS pH 7.4 and 0.1% Tween-20) and then blocked at room temperature for 1 hour. Next, 20 μL of the DNA handle construct (250 pM in SuperBlock PBS) was added to the flow cell and allowed to incubate at room temperature for 5 minutes before being washed with 80 μL of SuperBlock. Mono Mag Streptavidin Beads (4.5 μm, Ocean Nanotech) were prepared by washing twice with SuperBlock, then diluted to 500 μg/mL in SuperBlock. Then, 60 μL of beads were added to the flow cell and incubated for 2 minutes. The cell was washed again with 80 μL of SuperBlock, and finally, inlet and outlet ports were sealed with high vacuum grease.

### Validation: Test Run

The completed flow cell containing the bead-to-glass tethering was securely mounted in the sample chamber, and the objective was manually threaded into the objective focus tube and positioned adjacent to the coated glass slide. After the assembly was secured in the CFM microscope, a live image was brought up on screen using a video streaming client (VLC media player, VideoLAN Association) on the external controlling computer used for overall control. Wireless focusing on the tethered beads was accomplished by using the stepper motor connected to the rotating RPi to move the objective. A second focusing check was performed immediately after the CFM reached the desired experimental rotational velocity. Then, video streaming was stopped and Raspistil time-lapse image capture was initiated. WiFi-controlled adjustment of the microscope objective allows the user to refocus at any time during the experiment, such as when focus is compromised due to high forces. However, this was generally not necessary after the post-spinup focus was achieved.

For the test run, single-molecule force-dependent, DIG:α-DIG rupture events were monitored by taking snapshots every 5 s for 1430 s at 337 RPM and 478 RPM for 12 pN and 24 pN experiments, respectively. Images were manually processed in ImageJ (ImageJ.net) by filtering out the green channel to reduce background (see Figure 8). A dimensional threshold, including a minimum and maximum particle size was established to eliminate counting contaminants and/or bead aggregates. A total bead count per image was calculated using the “analyze particles” tool in XXXXX software (XXX). Bead counts were normalized to 1 based on the total number of beads in the first frame, and a rupture curve was plotted using Prism GraphPad for both 12 and 24 pN of force (see Figure 7).

## RESULTS

### Test Run Results

Two CFM test runs were performed at 337 RPM and 478 RPM, correlating to the calculated forces 12 pN and 24 pN, respectively. Force values were calculated by converting RPM values into angular velocity using Equation 2. The manufacturer provides the value for the buoyancy adjusted mass of the bead in the buffer solution (put the value here). Individual rupture events of the DIG:α-DIG interface were tracked at 5 s intervals over a 1430 s time course in each of the experiments. These data were normalized to the total number of beads present in the first frame. An example of a single frame of the bead count can be seen in Figure 8. In the 12 pN experiment, after about 1430 s, approximately 68% of beads remained in the frame, compared to the 24 pN run, where about 28% of beads remained.

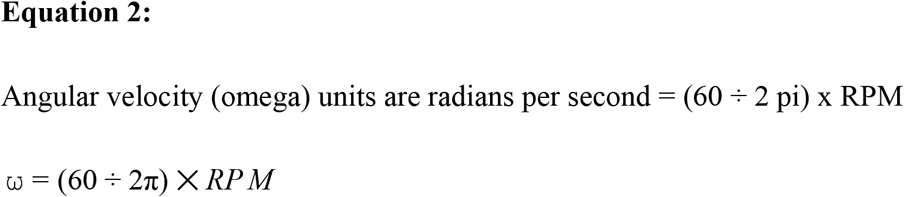

Both 12 pN and 24 pN data sets describing rupture event curves fit well to a one-phase decay model indicative of a homogeneous distribution of single-tethered beads (citation explaining how the curves can calculate forces, ie. the Y = plateau + (Yo - P)…). Because the streptavidin:biotin interaction is much stronger with respect to the DIG:α-DIG interaction (Neuert et al., 2006), rupture events correspond exclusively to the breaking of the DIG:α-DIG interaction (See Figure 5). The exponential decay of the experimental run at 12 pN indicates a half-life of 2664s, which was slower than the 24 pN exponential decay with a half-life of 518.8 s. These results suggest that rupture events are force-dependent and scale with the amount of force applied. More importantly, for the purpose of our validation testing, these experiments indicated that the CFM functioned effectively.

**Figure 7:**
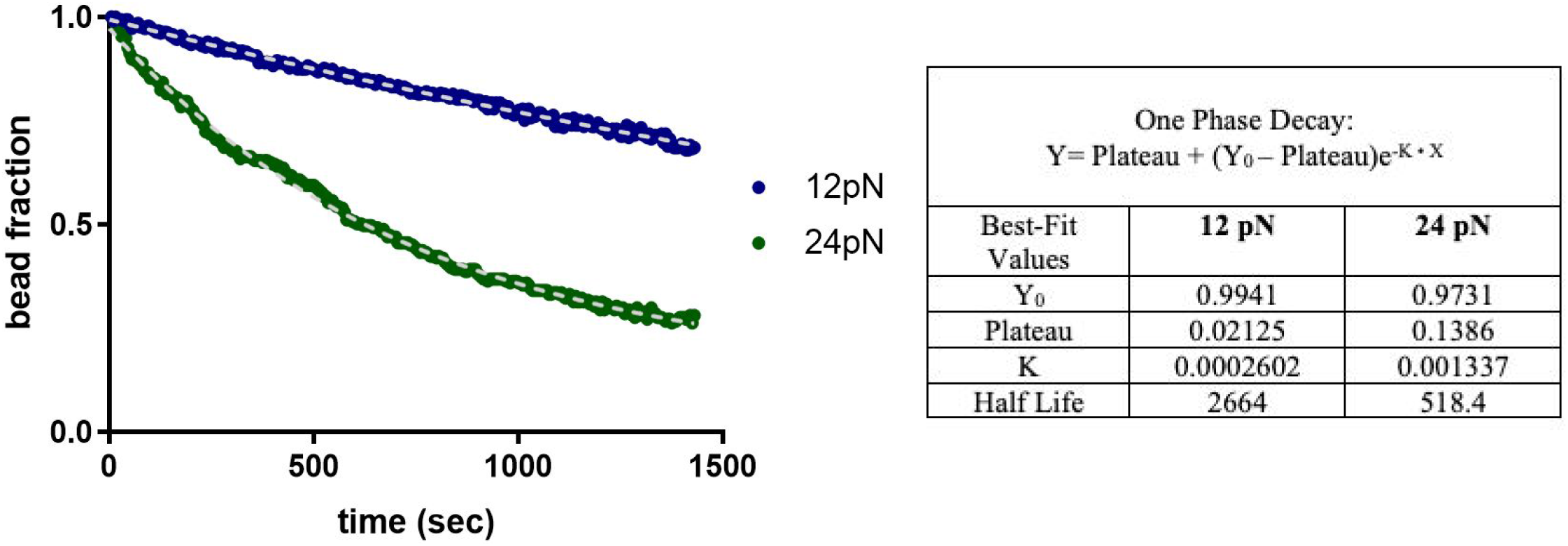
A scatter plot of single-molecule force-dependent DIG:αDIG normalized rupture events fit to a one phase decay (gray dashed lines) from two runs at calculated forces 12 pN and 24 pN, where Y_0_ is the y-intercept, Plateau is the y-asymptote, and K is the decay rate (s^-1^). The rupture half-life (s) is computed as ln(2)/K. The plots have a characteristic one-phase exponential decay where greater force correlates with more rupture events and a lower bead fraction over time.

**Figure 8:**
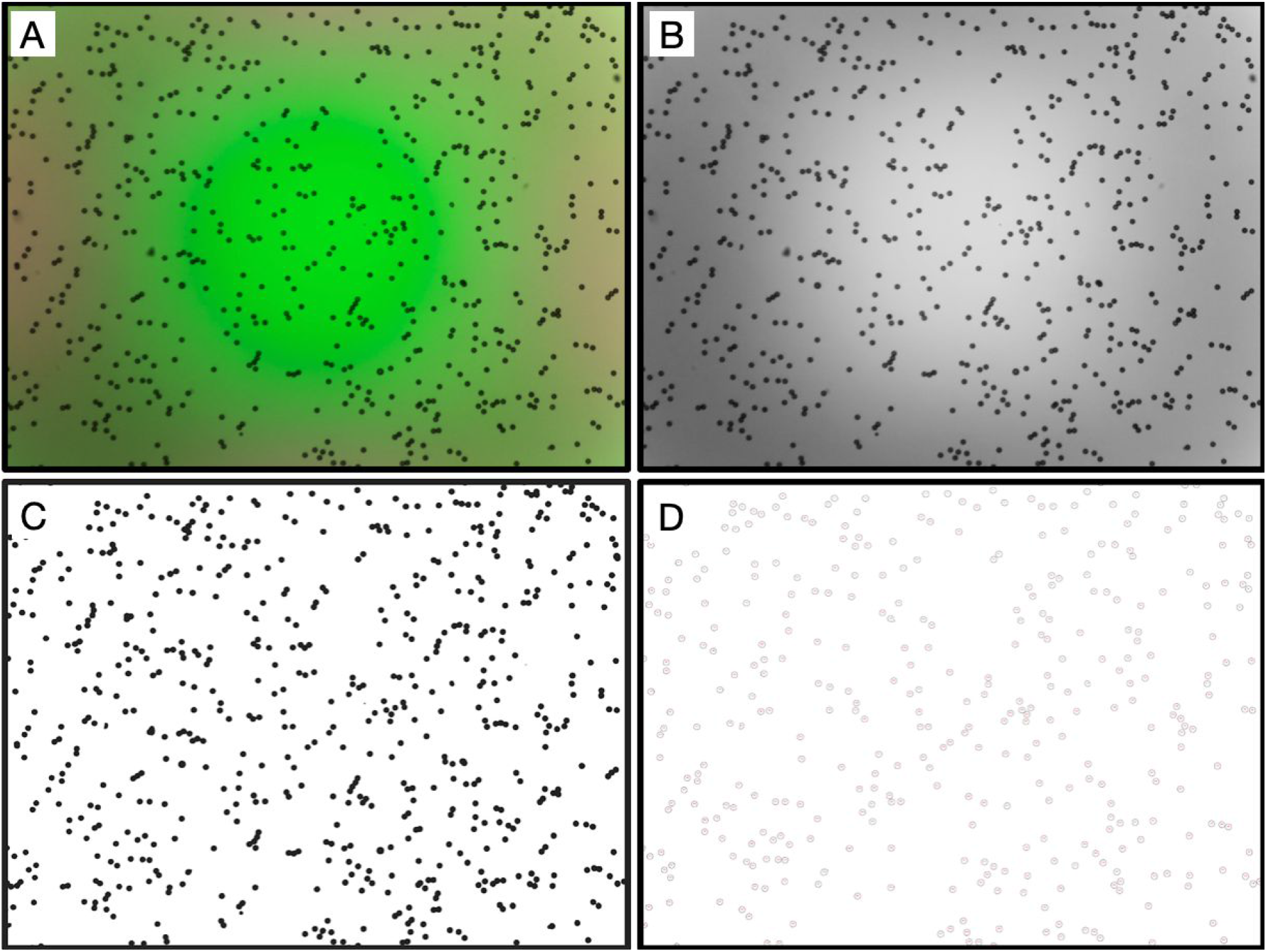
Example of image processing using ImageJ. The raw image (A) color channels are split and only the green channel (B) is selected. Then, background subtraction and thresholding is applied (C) leaving only single beads, which can be counted. Clusters of beads are excluded using a particle size cut-off (D).

### Educational Emphasis

The project’s overarching goal was not only to build a functional CFM, but also to create a learning environment where a team of students could collaborate and build a functional CFM. During CFM construction, students engaged in interdisciplinary collaboration, which strengthened their teaching and communication skills. In addition to learning from the students, the postgraduate researcher was able to develop his skills as an educator.

A qualitative survey was sent out to evaluate the research team’s experiences, receiving responses from 11 out of the 12 participants. The student responses were positive and indicated success in maintaining a collaborative project where student opinions and inputs were valued. Students also expressed appreciation for opportunities to acquire new skills with a wide range of applications, such as 3D printing and coding with Python. More detailed descriptions of the student experience are provided in the Student Quotes. Overall, the goal of producing a functional instrument with the collaboration of students from various educational backgrounds proved to be successful. Many individuals expressed a desire to work on future projects using their newly developed skills. The raw survey results are included in the supplemental information (S__).

### Student Quotes

“As a sophomore in high school, I didn’t have a large impact on the project, but it was very exciting to help out with 3D printer maintenance and part design for the research project. This was my first real experience in an “internship-like” setting, where I got to use the skills I had (and learn new ones) to work on a large project.” — high school student

“This project was extremely effective in promoting interaction with the research community and teaching you applicable career skills. I feel that I learned scientific research at a far deeper level than simply performing directed experiments for another researcher. Planning, performing, and analyzing experiments has given me the confidence and skills necessary to continue on as a career researcher.” — 4th year undergraduate student

“The main thing I learned is how insanely cool Gordon Lab’s research is. As I mentioned, this made me much more open to and enthusiastic about going into a natural-science-focused research area. Additionally, I improved my general programming abilities and learned to apply the skills I learned at my job as a web developer to new contexts.” — graduate student

Although the experience was positive overall, students provided suggestions for future improvements. Greater project organization was a common suggestion, which can easily be understood because the team consisted of many members. More specifically, students stated that they would have preferred to meet as a whole group to update members on the status of the different subprojects. Also, students said that they would have liked additional background information about the subprojects to make a more informed decision about the avenue they selected. These suggestions will be taken into account for future projects in hopes of further improving the research experience.

## DISCUSSION

### Evaluation of Design

The goal of designing and creating an operational, WiFi-enabled CFM was successful, both from the perspective of creating a useful instrument as well as doing so in an educationally edifying way. This CFM design incorporates (1) wireless communication for real-time control and data acquisition, (2) precise and programmable speed control, (3) real time microscope focusing, and (4) design modularity unconstrained by the size limitations of a commercial swinging-bucket centrifuge. The instrument imaged and analyzed force-dependent rupture events of macromolecular interactions under constant force using a modified version of a previous method of probing the DIG:α-DIG interaction (Yang et al., 2016).

Many groups have defined the force-dependent rupture kinetics of this interaction; unfortunately a wide range of force-dependent dissociation lifetimes have been reported over the years using various experimental set-ups and instrumentation resulting in conflicting results (Eeftens et al., 2015; Halvorsen & Wong, 2010; Neuert et al., 2006). Although CFMs are theoretically calibration-free instruments (Halvorsen & Wong, 2010), directly comparing experimental results to a calibrated MT instrument would validate rupture kinetics and permit the use of a broader range of forces, differing bead masses, and more replicates. With further testing, the presented CFM design and analysis set-up will be a powerful and accurate instrument to define force-dependent dissociation kinetics of biological macromolecules.

The use of smaller beads required higher spin speeds where dynamic unbalance became a limiting issue. Therefore, larger beads allow samples to be spun at lower speeds, mitigating the balance and safety issues. It is beneficial to spin briefly at a very low, standard RPM to clear non-specifically attached beads, preventing them from confounding the data set. Due to our design’s programmability, the clearing of non-specific beads can be incorporated into a data collection run that is easily reproducible. Other experimental designs that would take advantage of the ability to perform non-constant force experiments by varying the rotational speed can be developed.

### Educational Outcomes

This project’s development and success can be attributed to the team of determined and inquisitive high school and undergraduate students under the guidance of senior research mentors. Those typically in leadership roles, such as postdoctoral researchers and the graduate student first author, prioritized high school students and undergraduates in leadership roles during CFM design. Avenues of leadership included recruiting others to assist, teaching others as a way of learning, and mentoring others, all of which promoted collaboration in a research setting. Having the opportunity to work with mentors allowed students to develop their solutions within their subprojects. Due to the emphasis on student mentorship rather than direct instruction, individuals were better able to discover their strengths and interests.

Initially, bringing together students from different educational backgrounds was difficult due to a varying degree of knowledge on the subject matter. However, with the guidance of an experienced postdoctoral researcher and collaboration with others, students could bring their strengths to the subproject(s) of their choosing. Through working on these projects, students developed and refined their skills.

Overall, the goal of producing a final research project involving biochemistry, biophysics, and engineering in collaboration with students from various educational backgrounds proved to be successful. This project provided students with exposure to real-life applications of scientific concepts and techniques. However, for future projects, the organization will be improved with regular meetings to update the team on the project status and to provide more in-depth information on subprojects before students choose their focus area.

### Future Directions

The presented CFM design contains more available development opportunities than a swinging bucket CFM offers. The extra space provided by a purpose-built rotor design will allow for the incorporation of additional features in CFM V2, which will be discussed in detail below. In following this design route, we recognize that the development of a similar model using this paper because a guide is more difficult.

#### Addressing Dynamic Spin Unbalance

A design incorporating dual, radially symmetrical microscopes will allow experimental and control samples to be run simultaneously. Moreover, this radial symmetry will better address the problem of dynamic unbalance as equipment mass will be symmetrically distributed across the axis of rotation. Further, the introduction of radially-symmetric duplicates for each and every CFM component allows for the introduction of a hypergravity fluidics pumping system.

#### Biology

A major impetus for the development of inexpensive single-force spectroscopy instruments is to contribute to the knowledge of mechanoresponsive biological systems by expanding the ability of low-budget laboratories to do significant research in the field of mechanobiology. For example, the CFM V2 design will be able to characterize putative force-responsive biological proteins, such as the adhesion G-coupled protein receptor family (Scholz et al., 2016), with a reproducible, accurate, low-cost, and simple methodology. By equipping researchers with instruments to characterize mechanoresponsive biological systems, an improved understanding of microforces in biology can be developed.

#### Cell-Adhesion Assay Capabilities

The challenge of centrifuge-based cell detachment methods must be overcome in order to expand the versatility of the CFM in quantifying single-cell adhesive forces in real-time. (Giacomello et al., 1999; Reyes & García, 2003). Centrifuge Force Microscopy is used to quantify the force-dependent adherence at a single cell level of large populations of cells in response to culture environments or even drug treatments. Expanding this method so that it includes real-time visualization capabilities of an on-board microscope would enable real-time data acquisition abilities, higher measurement accuracy, and experimental design flexibility.

#### Safety

Vibration detection with comparison to nominal, leading to electronic braking and power shutoff.

Using plastics that are susceptible to degradation and are weaker than machined components.

#### Hypergravity Fluidics

Using the extra space available on the rotating plate of the CFM, a fluidics system that functions under a hypergravity situation can be introduced. This system will be composed of a stepper motor-actuated pump system that can provide precision, biological, buffer solution delivery through the flow cell while spinning. The movement of a buffer solution from one syringe to another can present problems with balance. However, the use of two fluidics systems that are positioned on the rotor with radial symmetry and would operate in tandem, each moving the exact same quantities of fluid, would mitigate unbalance. Further, the use of a second microscope will allow for simultaneous runs of experimental and control systems.

#### Use of Commercially-Available Components and Alternative Plastics

A preference for the use of off-the-shelf parts in version 2, as opposed to custom-machined or 3D-printed parts used in the presented CFM, will allow the design to be more easily constructed by other labs. The PLA plastic used for all of the printer components in version 1 performed adequately. However, these parts are more prone to vibration-inducing flexing and stress failure over time when compared to high precision, machined-metal pieces available from suppliers such as Thorlabs. Alternative plastics, such as ABS and PETG, are also being considered and tested for CFM V2 when purchasable parts are not available. Rigid plastics (PLA) that do not flex under spinning forces are preferable compared to plastics that bend and flex easily (ABS and PETG). This preference is due to the potential of ABS and PETG plastics to cause unbalance issues, however, strength is also a consideration. While material strength and vibration considerations were not addressed with the current version discussed herein, CFM V2 will be designed and tested with these aspects in mind.

## Supplemental Info Google Doc Link

https://docs.google.com/document/d/11AOQVZPpu4Q8flva4qkYTx201XiCdJ_NKPDabhv7NTM/edit

## Acknowledgements

We wish to thank William (Bill) Voje and Michael Rother of the UMN Student Physics Shop for their help in machining and in training our team to do machining. We could not have built this CFM without them.

